# Extensive Copy Number Variation in Fermentation-Related Genes among *Saccharomyces cerevisiae* Wine Strains

**DOI:** 10.1101/105502

**Authors:** Jacob Steenwyk, Antonis Rokas

## Abstract

Due to the importance of *Saccharomyces cerevisiae* in wine-making, the genomic variation of wine yeast strains has been extensively studied. One of the major insights stemming from these studies is that wine yeast strains harbor low levels of genetic diversity in the form of single nucleotide polymorphisms (SNPs). Genomic structural variants, such as copy number (CN) variants, are another major type of variation segregating in natural populations. To test whether genetic diversity in CN variation is also low across wine yeast strains, we examined genome-wide levels of CN variation in 132 whole-genome sequences of *S. cerevisiae* wine strains. We found an average of 97.8 CN variable regions (CNVRs) affecting ~4% of the genome per strain. Using two different measures of CN diversity, we found that gene families involved in fermentation-related processes such as copper resistance (*CUP*), flocculation (*FLO*), and glucose metabolism (*HXT*), as well as the *SNO* gene family whose members are expressed before or during the diauxic shift showed substantial CN diversity across the 132 strains examined. Importantly, these same gene families have been shown, through comparative transcriptomic and functional assays, to be associated with adaptation to the wine fermentation environment. Our results suggest that CN variation is a substantial contributor to the genomic diversity of wine yeast strains and identify several candidate loci whose levels of CN variation may affect the adaptation and performance of wine yeast strains during fermentation.

## Introduction

*Saccharomyces cerevisiae*, commonly known as baker’s or brewer’s yeast, has been utilized by humans for the production of fermented beverages since at least 1,350 B.C.E. but may go as far back as the Neolithic period 7,000 years ago (Mortimer 2000; Cavalieri *et al*. 2003). Phylogenetic analyses and archaeological evidence suggest wine strains originated from Mesopotamia (Bisson 2012) and were domesticated in a single event around the same time as the domestication of grapes (Schacherer *et al*. 2009; Sicard and Legras 2011). Further phylogenetic, population structure and identity-by-state analyses of single nucleotide polymorphism (SNP) data reveal close affinity and low genetic diversity among wine yeast strains across the globe, consistent with a domestication-driven population bottleneck (Liti *et al*. 2009; Schacherer *et al*. 2009; Sicard and Legras 2011; Cromie *et al*. 2013; Borneman *et al*. 2016). These low levels of genetic diversity have led some to suggest that further wine strain development should be focused on introducing new variation into wine yeasts rather than exploiting their standing variation (Borneman *et al*. 2016).

Many wine strains have characteristic variants that have presumably been favored in the wine-making environment (Marsit and Dequin 2015). For example, adaptive point mutations, deletions and rearrangements in the promoter and coding sequence of *FLO11* contribute to flocculation and floating thereby increasing yeast cells’ ability to obtain oxygen in the hypoxic environment of liquid fermentations (Fidalgo *et al*. 2006). Similarly, duplications of *CUP1* are strongly associated with resistance to copper (Warringer *et al*. 2011), which at high concentrations can cause stuck fermentations, and *THI5*, a gene involved in thiamine metabolism whose expression is associated with an undesirable rotten-egg sensory perception in wine, is absent or down regulated among wine strains and their derivatives (Bartra *et al*. 2010; Brion *et al*. 2014). As these examples illustrate, the mutations underlying these, as well as many other, presumably adaptive traits are not only single nucleotide polymorphisms (SNPs), but also genomic structural variants, such as duplications, insertions, inversions, and translocations (Pretorius 2000; Marsit and Dequin 2015).

Copy number (CN) variants, a class of structural variants defined as duplicated or deleted loci ranging from 50 bp to whole chromosomes (Zhang *et al*. 2009; Arlt *et al*. 2014), have recently started receiving considerable attention due to their widespread occurrence (Sudmant *et al*. 2010; Bickhart *et al*. 2012; Axelsson *et al*. 2013; Pezer *et al*. 2015) as well as their influence on gene expression and phenotypic diversity (Freeman *et al*. 2006; Henrichsen *et al*. 2009). Mechanisms of CN variant evolution include non-allelic homologous recombination (Lupski and Stankiewicz 2005) and retrotransposition (Kaessmann *et al*. 2009). CN variants are well studied in various mammals, including humans (*Homo sapiens*; Sudmant *et al*. 2015), cattle (*Bos taurus*; Bickhart *et al*. 2012), the house mouse (*Mus musculus*; Pezer *et al*. 2015), and the domestic dog (*Canis lupus familiaris*; Axelsson *et al*. 2013), where they are important contributors to genetic and phenotypic diversity.

Relatively few studies have investigated whole-genome CN profiles in fungi (Hu *et al*. 2011; Farrer *et al*. 2013; Steenwyk *et al*. 2016). For example, the observed CN variation of chromosome 1 in the human pathogen *Cryptococcus neoformans* results in the duplications of *ERG11*, a lanosterol-14-α-demethylase and target of the triazole antifungal drug fluconazole (Lupetti *et al*. 2002), and *AFR1*, an ATP binding cassette (ABC) transporter (Sanguinetti *et al*. 2006), leading to increased fluconazole resistance (Sionov *et al*. 2010). Similarly, resistance to itraconazole, a triazole antifungal drug, is attributed to the duplication of cytochrome P-450-depdendent C-14 lanosterol α-demethylase (*pdmA*) – a gene whose product is essential for ergosterol biosynthesis – in the human pathogen *Aspergillus fumigatus* (Osherov *et al*. 2001). Finally, in the animal pathogen *Batrachochytrium dendrobatidis*, the duplication of Supercontig V is associated with increased fitness in the presence of resistance to an antimicrobial peptide, although the underlying genetic elements involved remain elusive (Farrer *et al*. 2013).

Similarly understudied is the contribution of CN variation to fungal domestication (Gibbons and Rinker 2015; Gallone *et al*. 2016). Notable examples of gene duplication being associated with microbial domestication include those of α-amylase in *Aspergillus oryzae*, which is instrumental in starch saccharification during the production of sake (Hunter *et al*. 2011; Gibbons *et al*. 2012), and of the *MAL1* and *MAL3* loci in beer associated strains of *S. cerevisiae*, which metabolize maltose, the most abundant sugar in the beer wort (Gallone *et al*. 2016; Gonçalves *et al*. 2016). Beer strains of *S. cerevisiae* often contain additional duplicated genes associated with maltose metabolism, including *MPH2* and *MPH3*, two maltose permeases, and the putative maltose-responsive transcription factor, *YPR196W* (Gonçalves *et al*. 2016). Adaptive gene duplication in *S. cerevisiae* has also been detected in experimentally evolved populations (Dunham *et al*. 2002; Gresham *et al*. 2008; Dunn *et al*. 2012). Specifically, duplication of the locus containing the high affinity glucose transporters *HXT6* and *HXT7* has been observed in adaptively evolved asexual strains (Kao and Sherlock 2008) as well as in populations grown in a glucose-limited environment (Brown *et al*. 1998; Dunham *et al*. 2002; Gresham *et al*. 2008). Altogether, these studies suggest that CN variation is a significant contributor to *S. cerevisiae* evolution and adaptation.

To determine the contribution of CN variation to genome evolution in wine strains of *S. cerevisiae*, we characterized patterns of CN variation across the genomes of 132 wine strains and determined the functional impact of CN variable genes in environments reflective of wine-making. Our results suggest that there is substantial CN variation among wine yeast strains, including in gene families (such as *CUP, FLO, HXT* and *MAL*) known to be associated with adaptation in the fermentation environment. More generally, it raises the hypothesis that CN variation is an important contributor to adaptation during microbial domestication.

## Methods

### Data Mining, Quality Control and Mapping

Raw sequence data for 132 *Saccharomyces cerevisiae* wine strains were obtained from three studies (Borneman *et al*. 2016, 127 strains, Bioproject ID: PRJNA303109; Dunn *et al*. 2012, 2 strains, Bioproject ID: SRA049752; Skelly *et al*. 2013, 3 strains, Bioproject ID: PRJNA186707) (Figure S1, File S1). Altogether, these 132 strains represent a diverse set of commercial and non-commercial isolates from the ‘wine’ yeast clade (Borneman *et al*. 2016).

Sequence reads were quality-trimmed using TRIMMOMATIC, version 0.36 (Bolger *et al*. 2014) with the following parameters and values: leading:10, trailing:10, slidingwindow:4:20, minlen:50. Reads were then mapped to the genome sequence of the *S. cerevisiae* strain S288c (annotation release: R64.2.1; http://www.yeastgenome.org/) using BOWTIE2, version 1.1.2 (Langmead and Salzberg 2012) with the ‘sensitive’ parameter on. For each sample, mapped reads were converted to the bam format, sorted and merged using SAMTOOLS, version 1.3.1. Sample depth of coverage was obtained using the SAMTOOLS depth function (Li *et al*. 2009).

### CN Variant Identification

To facilitate the identification of single nucleotide polymorphisms (SNPs), we first generated mpileup files for each strain using SAMTOOLS, version 1.3.1 (Li *et al*. 2009). Using the mpileup files as input to VARSCAN, version 2.3.9 (Koboldt *et al*. 2009, 2012), we next identified all statistically significant SNPs (Fisher’s Exact test; *p* < 0.05) present in the 132 strains that had a read frequency of at least 0.75 and minimum coverage of 8X. This step enabled us to identify 149,782 SNPs. By considering only SNPs that harbored a minor allele frequency of at least 10%, we retained 43,370 SNPs. These SNPs were used to confirm the evolutionary relationships among the strains using Neighbor-Net phylogenetic network analyses in SPLITSTREE, version 4.14.1 (Huson 1998) as well as the previously reported low levels of SNP diversity (Figure S2; Borneman *et al*. 2016).

To detect and quantify CN variants we used CONTROL-FREEC, version 9.1 (Boeva *et al*. 2011, 2012), which we chose because of its low false positive rate and high true positive rate (Duan *et al*. 2013). Importantly, the average depth of coverage or read depth of the 132 strains was 30.1 ± 14.7X (minimum: 13.0X, maximum: 104.5X; Figure S3), which is considered sufficient for robust CNV calling (Sims *et al*. 2014).

CONTROL-FREEC uses LOESS modeling for GC-bias correction and a LASSO-based algorithm for segmentation. Implemented CONTROL-FREEC parameters included window = 250, minExpectedGC = 0.35, maxExpectedGC = 0.55 and telocentromeric = 7000. To identify statistically significant CN variable loci (*p* < 0.05), we used the Wilcoxon Rank Sum test. The same CONTROL-FREEC parameters, but with a window size of 25 base pairs (bp), were used to examine CN variation within the intragenic Serine/Threonine-rich sequences of *FLO11* (Lo and Dranginis 1996). BEDTOOLS, version 2.25 (Quinlan and Hall 2010) was used to identify duplicated or deleted genic loci (i.e., CN variable loci) that overlapped with genes by at least one nucleotide. The CN of each gene (genic CN) was then calculated as the average CN of the 250 bp windows that overlapped with the gene’s location coordinates in the genome. The same method was used to determine non-genic CN for loci that did not overlap with genes (ie., non-genic CN variable loci). To identify statistically significant differences between CN variable loci that were duplicated versus those that were deleted, we employed the Mann-Whitney *U* test (Wilcoxon rank-sum test) with continuity correction (Wallace 2004).

### Diversity in CN Variation and GO Enrichment

To identify CN diverse loci we used two different measures. The first measure calculates the statistical variance (s^2^) for each locus where CN variants were identified in one or more strains. s^2^ values were subsequently log_10_ normalized. Log_10_(s^2^) accounts for diversity in raw CN values but not for diversity in CN allele frequencies. Thus, we also employed a second measure based on the Polymorphic Index Content (PIC) algorithm, which has previously been used to identify informative microsatellite markers for linkage analyses by taking into account both the number of alleles present and their frequencies (Keith *et al*. 1990; Risch 1990). PIC has also been used to quantify population-level diversity of simple sequence repeat loci and restriction fragment length polymorphisms in maize (Smith *et al*. 1997). PIC values were calculated for each locus harboring at least one CN variant based on the following formula:

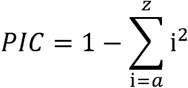

where i^2^ is the squared frequency of *a* to *z* CN values (Smith *et al*. 1997). PIC values may range from 0 (no CN diversity) to 1 (all CN alleles are unique).

To create a list of loci exhibiting high CN diversity for downstream analyses, we retained only those loci that fell within the 50^th^ percentile of log_10_(s^2^) values (min = −2.12, median = −1.02, and max = 2.40) or the 50^th^ percentile of PIC values (min = 0.02, median = 0.14, and max = 0.96).

Genes overlapping with loci exhibiting high CN diversity were used for Gene ontology (GO) enrichment analysis with AMIGO2, version 2.4.24 (Carbon *et al*. 2009) using the PANTHER Overrepresentation Test (release 20160715) with default settings. This test uses the PANTHER Gene Ontology database, version 11.0 (Thomas *et al*. 2003; release date 2016-07-15) which is directly imported from the GO Ontology database, version 1.2 (GeneOntologyConsortium 2004; release date 2016-10-27), a reference gene list from *S. cerevisiae*, and a Mann-Whitney *U* test (Wilcoxon rank-sum test) with Bonferroni multi-test corrected *p*-values to identify over-and under-represented GO terms (Mi *et al*. 2013). Statistical analyses and figures were created using PHEATMAP, version 1.0.8 (Kolde 2012), GPLOTS, version 3.0.1, GGPLOT2 (Wickham 2009) or standard functions in R, version 3.2.2 (R Development Core Team 2011).

### Identifying Loci Absent in the Reference Strain

To identify loci absent from the reference strain but present in other strains, we assembled unmapped reads from the 20 strains with the lowest percentage of mapped reads. The percentage of mapped reads was determined using SAMTOOLS (Li *et al*. 2009); its average across strains was 96% (min = 70.5% and max = 99%; Figure S4). Unmapped reads from the 20 strains with the lowest percentage of mapped reads were assembled using SPADES, version 3.8.1 (Bankevich *et al*. 2012). The identity of scaffolds longer than the average length of a *S. cerevisiae’s* gene (~1,400 bp) was determined using blastx from NCBI’s BLAST, version 2.3.0 (Madden 2013) against a local copy of the GenBank non-redundant protein database (downloaded on January 5, 2017).

## Results

### Descriptive Statistics of CN variation

To examine CN variation across wine yeasts, we generated whole genome CN profiles for 132 strains (Figure S5, File S2). Across all strains, we identified a total of 2,820 CNVRs that overlapped with 2,061 genes and spanned 3.7 megabases (Mb). The size distribution of CNVRs was skewed toward CN variants that were shorter than 1 kilobase (kb) in length (Figure 1A, Figure S6A & Table S1). Strains had an average of 97.8 ± 9.5 CNVRs (median = 86) (Figure S6B) that affected an average of 4.3% ± 0.1% of the genome (median = 4.1%) (Figure S6C).

**Figure 1.**
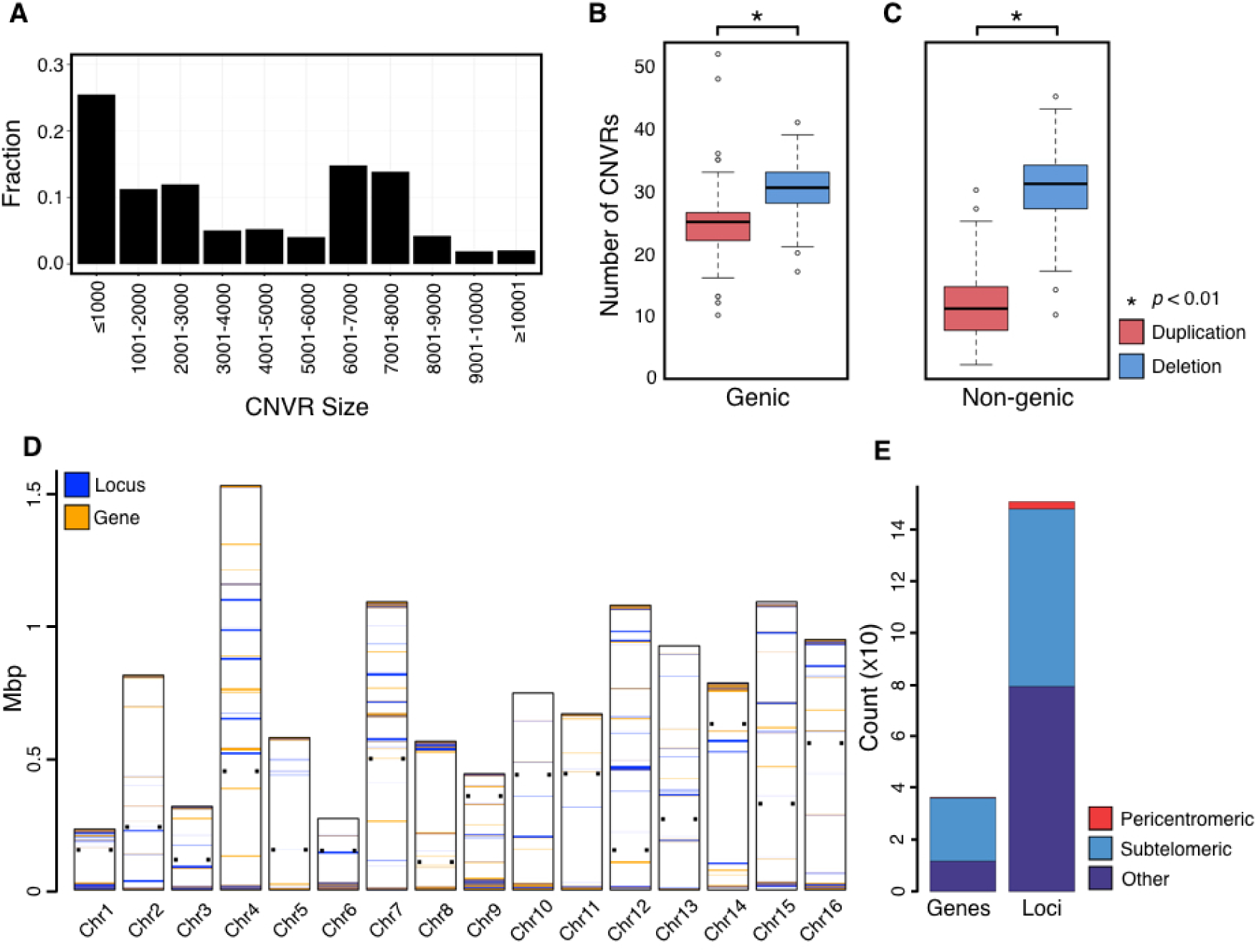
**Size distribution and location of CN variable loci**. (A) The fraction of CN variable regions (CNVRs) (y-axis) for a given size range. Most CNVRs are less than or equal to 1,000bp. (B, C) Deleted genic (B) and non-genic (C) CNVRs are more prevalent than duplicated ones (*p* < 0.01 for both comparisons). (D) Location of CN variable loci across the 16 yeast chromosomes. The small, black squares on either side of each chromosome denote centromere location. Loci (blue bars) and genes (orange bars) harboring high log_10_(s^2^) or PIC values are shown. (E) 684 of the 1,502 CN diverse loci and 243 of the 363 CN diverse genes reside in subtelomeric regions of the yeast genome; in contrast, very few are found in pericentromeric regions (28 loci and 3 genes).

Due to the known influence of CN variable genes (Henrichsen *et al*. 2009; Orozco *et al*. 2009), we next quantified the number of genic and non-genic CNVRs (Figure 1B and C). We found statistically significant differences in the number of duplicated and deleted loci that are genic or non-genic (Mann-Whitney U test; *p* < 0.01 for both genic and non-genic comparisons) revealing that there were significantly more deleted genic and non-genic CNVRs than duplicated ones.

### CN Diversity in Subtelomeres

To identify loci that exhibited high CN diversity, we retained only those loci that fell within the 50^th^ percentile of at least one of our two different measures log_10_(s^2^) and PIC) across the 132 strains. The distributions of the two measures (Figure S7) were similar, with 1,326 loci (Figure S7C) and 291 genes (Figure S7D) identified in the top 50% of CN diverse genes by both measures.

In addition, the log_10_(s^2^) measure identified an additional 85 loci and 54 genes in its set of top 50% genes, and PIC an additional 85 loci and 18 genes. In total, our analyses identified 1,502 loci and 363 genes showing high CN diversity. Among the genes harboring the highest log_10_(s^2^) and PIC values were *YLR154C-G* (PIC = 0.96; log_10_(s^2^) = 2.16), *YLR154W-A* (PIC = 0.96; log_10_(s^2^) = 2.16), *YLR154W-B* (PIC= 0.96; log_10_(s^2^) = 2.16), *YLR154W-C* (PIC = 0.96; log_10_(s^2^) = 2.16), *YLR154W-E* (PIC = 0.96; log_10_(s^2^) = 2.16),*YLR154W-F* (PIC = 0.96; log_10_(s^2^) = 2.16) and *YLR154C-H* (PIC = 0.93; log_10_(s^2^) = 2.40); these genes are all encoded within the 25S rDNA or 35S rDNA locus. The rDNA locus is known to be highly CN diverse (Gibbons *et al*. 2015) thereby demonstrating the utility and efficacy of our CN calling protocol as well as our two measures of CN diversity. We next generated CN diversity maps for all 16 *S.cerevisiae* chromosomes (Figure 1D; Figure S8). CN diversity was higher in loci and genes located in subtelomeres (defined as the 25 kb of DNA immediately adjacent to the chromosome ends; Barton *et al*. 2003). Specifically, 684 / 1,502 (45.5%) of CN diverse loci and 243 / 363 (66.9%) CN diverse genes were located in the subtelomeric regions. Conducting the same analysis using an alternative definition of subtelomere (defined as the DNA between the chromosome’s end to the first essential gene (Winzeler *et al*. 1999)) showed similar results. Specifically, 721 / 1,502 (48%) of CN diverse loci and 233 / 363 (64.2%) of CN diverse genes were located in the subtelomeric regions.

### GO Enrichment of CN Diverse Genes

To determine the functional categories over-and under-represented in the 363 genes showing high CN diversity, we performed GO enrichment analysis. The majority of enriched GO terms were associated with metabolic functions such as α-GLUCOSIDASE ACTIVITY (*p* < 0.01) and CARBOHYDRATE TRANSPORTER ACTIVITY (*p* < 0.01) (Figure 2 and File S3).

Genes associated with these GO terms include *SUC2* (*YIL162W*, involved in hydrolyzing sucrose), all six members from the *MAL* gene family (involved in the fermentation of maltose and other carbohydrates) and all five members of the *IMA* gene family (involved in isomaltose, sucrose and turanose metabolism). Other enriched categories were associated with multi-cellular processes such as the FLOCCULATION (*p* < 0.01) and AGGREGATION OF UNICELLULAR ORGANISMS (*p* = 0.03). All members of the *FLO* gene family (involved in flocculation) and *YHR213W* (a flocculin-like gene) were associated with these GO enriched terms.

**Figure 2.**
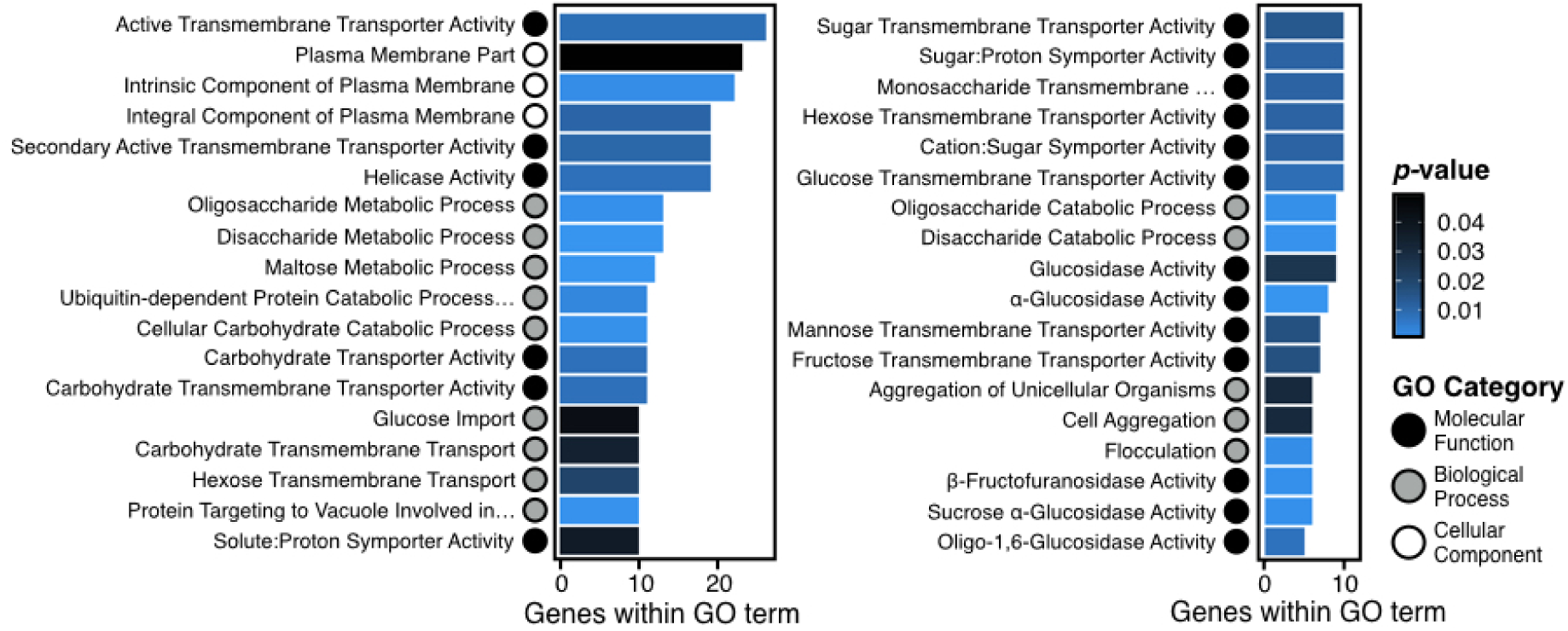
**Gene Ontology enriched terms from high CN diverse genes.** Molecular function (black), biological process (grey) and cellular component (white) GO categories are represented by circles and are enriched among the 363 genes that overlap with CN diverse loci. Enriched terms are primarily related to metabolic function, such as α-GLUCOSIDASE ACTIVITY (*p <* 0.01), CARBOHYDRATE TRANSPORTER ACTIVITY (*p <* 0.01) and FLOCCULATION (*p* < 0.01).

Contrary to overrepresented GO terms, underrepresented terms were associated with genes whose protein products are part of the interactome or protein-protein interactions such as PROTEIN COMPLEX (*p* < 0.01), MACROMOLECULAR COMPLEX ASSEMBLY (*p* = 0.03), TRANSFERASE COMPLEX (*p* < 0.01) and RIBONUCLEOPROTEIN COMPLEX BIOGENESIS (*p* = 0.04). Our finding of underrepresented GO terms being associated with multi-unit protein complexes supports the gene balance hypothesis, which states that the stoichiometry of genes contributing to multi-subunit complexes must be maintained to conserve kinetics and assembly properties (Birchler and Veitia 2010, 2012). Thus, genes associated with multi-unit protein complexes are unlikely to exhibit CN variation.

### Genic CN Diversity

To further understand the structure of CN variation in highly diverse CN genes, we first calculated the absolute CN of 23 genes associated with GO enriched terms related to wine fermentation processes (e.g., metabolic functions; Figure 2 and File S3) as well as 57 genes with the highest PIC or log_10_(s^2^) values (Figure S9 and File S4; 69 total unique genes). Among these 69 genes, gene CN ranged from 0 to 92; both the highest CN diversity and absolute CN values were observed in segments of the rDNA locus (mentioned above).

Importantly, 35 of the 69 genes have also been reported to have functional roles in fermentation-related processes. For example, the CNs of *PAU3* (*YCR104W*), a gene active during alcoholic fermentation, and its gene neighbor *ADH7* (*YCR105W*), an alcohol dehydrogenase, both varied between 0 and 3. Similarly, the absolute CN of the locus containing both *CUP1-1* (*YHR053C*; PIC = 0.868) and its paralog *CUP1-2* (*YHR055C*; PIC = 0.879) ranged from 0-14 (Figure 3; File S4), with 90 strains (68.2%) showing duplications (i.e., a CN greater than 1) and another 11 strains (8.3%) a deletion (i.e., a CN of 0). Interestingly, multiple copies of *CUP1* confer copper resistance to wine strains of *S. cerevisiae*, with CN variation at this locus thought to be associated with domestication (Warringer *et al*. 2011; Marsit and Dequin 2015).

**Figure 3.**
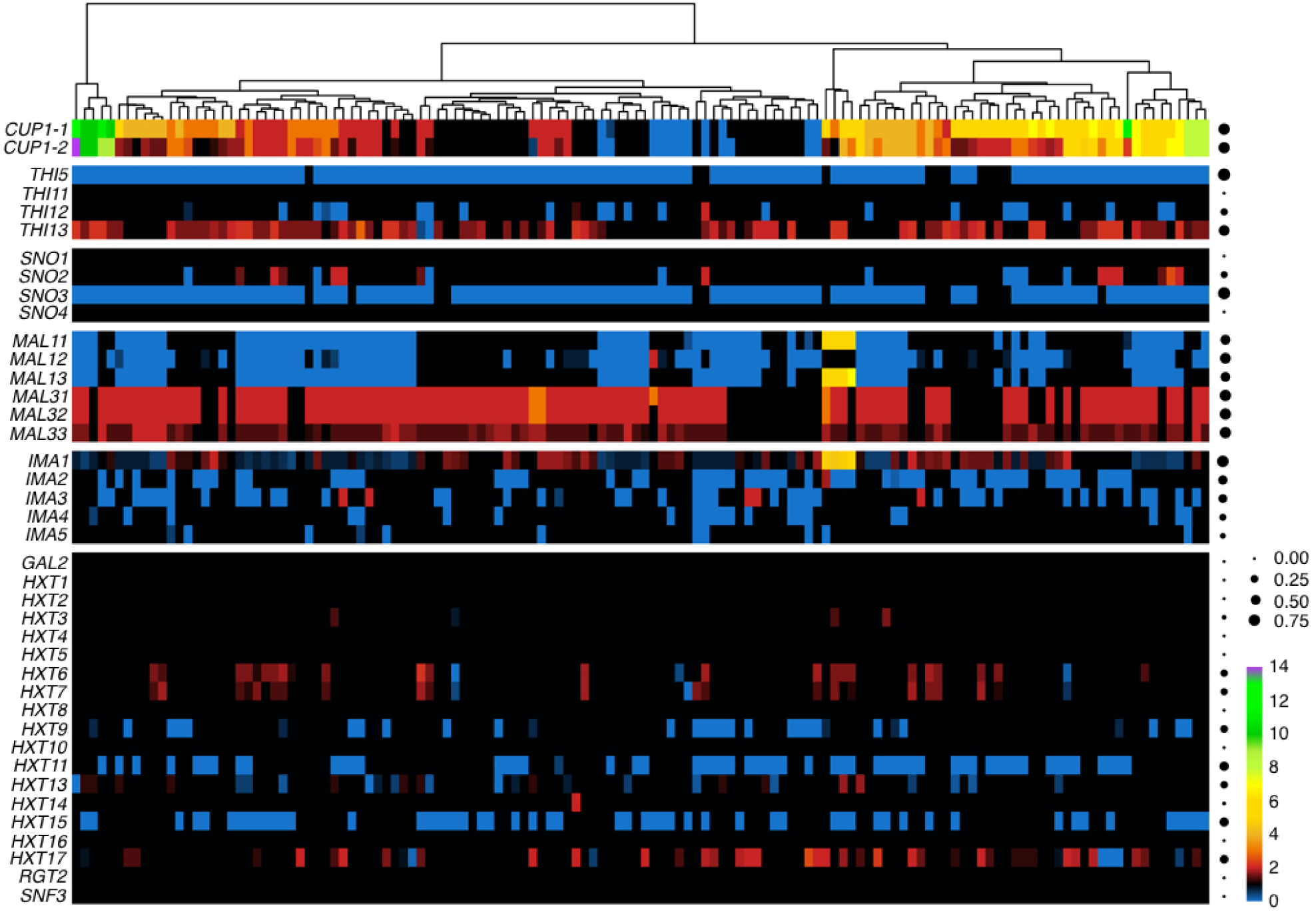
**CN variation of genes and gene families.** Heat map of the CN profiles the*CUP*, *THI*, *SNO*, *MAL*, *IMA* and *HXT* gene families; rows correspond to genes and columns to strains. Blue-colored cells correspond to deletions, black-colored cells to no CN variation and red-to-purple-colored cells to duplications (ranging from 2-14). Dots on the right side of the figure represent the proportion of individual strains that harbor CN variation in that gene - the larger the dot, the greater the proportion of the strains that is CN variable for that gene.

The expression of *SNO* family members is induced just prior to or after the diauxic shift as a response to nutrient limitation and is associated with vitamin B acquisition (Padilla *et al*. 1998; Rodríguez-Navarro *et al*. 2002). We found that *SNO2* (*YNL334C*) and *SNO3* (*YFL060C*) were among the 363 genes with highest CN diversity. *SNO2* was duplicated in 14 strains (10.6%) and deleted in 9 strains (6.8%), while *SNO3* was deleted in 117 strains (88.6%) (Figure 3). The other two members of the SNO gene family, *SNO1* (*YMR095C*) and *SNO4* (*YMR322C*), both showed a CN of 1 in all strains.

Another gene family whose members show high CN diversity is the *THI* gene family, which is responsible for thiamine metabolism and is activated at the end of the growth phase during fermentation (Brion *et al*. 2014). Specifically, *THI13* (*YDL244W;* PIC = 0.759) was among the 57 genes with the highest CN diversity (File S4), and *THI5* (*YFL058W*) and *THI12* (*YNL332W*) among the 363 most CN diverse genes (File S3). *THI13* was duplicated in 82 strains (62.1%) and deleted in 2 strains (1.5%) (Figure 3). In contrast, *THI5* was deleted in 121 strains (91.67%), whereas *THI12* was deleted in 23 strains (17.42%) and duplicated in only 3 strains (2.27%). Lastly, the CN of the last *THI* gene family member, *THI11* (*YJR156C*), did not exhibit CN variation.

In addition to the high CN diversity observed in all six members of the *MAL1* and *MAL3* loci responsible for maltose metabolism and growth on sucrose (Stambuk *et al*. 2000; Gallone *et al*. 2016), *MAL13* (*YGR288W*; PIC = 0.53) was among the 57 genes with the highest CN diversity (File S4). Evaluation of the absolute CN of all *MAL1* locus genes (Figure 3) showed that *MAL11, MAL12* (*YGR292W*), and *MAL13* were deleted in 65 (49.2%), 86 (65.2%), and 61 strains (46.2%), respectively. In contrast, the *MAL3* locus genes *MAL31* (*YBR298C*), *MAL32* (*YBR299W*), and *MAL33* (*YBR297W*) were duplicated in 100 (75.8%), 99 (75%), and 98 strains (74.2%), respectively. Interestingly, we did not observe any deletions in any of the *MAL3* locus genes across the 132 strains. When considering all members of the *MAL* gene family, we found that the 132 strains differed widely in their degree to which the locus had undergone expansion or contraction (Figure S10).

All members of the *IMA* gene family, composed of genes aiding in sugar fermentation (Teste *et al*. 2010), were among the 363 genes with high CN diversity (File S3) and *IMA1* (*YGR287C*; PIC = 0.87) was among the top 57 genes with the highest CN diversity (Files S4). *IMA1* was deleted in 54 strains (40.9%) and duplicated in 50 strains (37.9%) (Figure 3). Although many duplications or deletions did not span the entirety of *IMA1*, there were 4 strains that harbored high CNs between 4 and 6. These same four strains also had similar and unique duplications of *MAL11* and *MAL13*, suggesting that *IMA1*, *MAL11*, and *MAL13*, which are adjacent to each other in the genome, may have been duplicated as one locus. The other isomaltases (*IMA2-5*; *YOL157C*, *YIL172C*, *YJL221C* and *YJL216C*) were deleted in at least 11 strains (8.3%) and at most 55 strains (41.7%). No duplications in *IMA2*-5 were detected and only rarely in *IMA3* (5 strains, 3.8%). Altogether, the 132 strains exhibited both expansions and contractions of the *IMA* gene family (Figure S10).

We identified 7 members of the *HXT* gene family (*HXT6*/*YDR343C*, *HXT7*/*YDR342C*, *HXT9/YJL219W*, *HXT11/YOL156W*, *HXT13/YEL069C*, *HXT15*/*YDL245C*, and *HXT17*/*YNR072W*), which is involved in sugar transport, that were among the 363 CN diverse genes (File S3). Members of the *HXT* gene family were duplicated, deleted or had mosaic absolute CN values across the 132 strains. For example, *HXT6* and *HXT7* were primarily duplicated in 25 (18.9%) and 22 strains (16.7%), respectively, while only 3 strains (2.3%) had deletions in either gene (Figure 3). *HXT9*, *HXT11*, *HXT15* were deleted in 32 (24.2%), 57 (43.2%) and 53 strains (40.2%), respectively, while no strains had duplications. Finally, *HXT13* was duplicated in 12 strains (9.1%) and deleted in 17 strains (12.9%), and *HXT17* was duplicated in 37 strains (28%) and deleted in 9 strains (6.8%).

As expansions in the *HXT* gene family are positively correlated with aerobic fermentation in *Saccharomyces paradoxus* and *S. cerevisiae* (Lin and Li 2011), we also examined the absolute CN of all other 10 members (*GAL2/YLR081W, HXT1/YHR094C, HXT2/YMR011W, HXT4/YHR092C, HXT5/YHR096C, HXT8/YJL214W, HXT10/YFL011W, HXT16/YJR158W, RGT2/YDL138W*, and *SNF3/YDL194W*) of the *HXT* gene family (Figure 3). Interestingly, all remaining 10 members of the *HXT* gene family were not CN variable. Altogether, examination of the *HXT* family CN diversity patterns across the 132 strains suggests that wine yeast strains typically exhibit minor contractions (i.e., *HXT* gene deletions exceed those of duplications) relative to the S288c reference strain (Figure S10).

All five members of the *FLO* gene family, which is responsible for flocculation (Govender *et al*. 2008), a trait shown to aid in the escape of oxygen limited environments during liquid fermentation (Fidalgo *et al*. 2006; Govender *et al*. 2008), were found to be among the 363 most CN diverse genes. Furthermore, *FLO5* (*YHR211W*; PIC = 0.82) and *FLO11* (*YIR019C*; PIC = 0.88) were among the 57 genes with the highest CN diversity (File S4). Due to the importance of site directed CN variation in *FLO* family genes (Fidalgo *et al*. 2006), we modified our representation of CN variation to display intragenic CN variation using a 250 bp window (Figure S11). *FLO5* was partially duplicated in 57 strains (43.2%), partially deleted in 47 strains (35.6%) and 115 strains (87.1%) had at least one region of the gene unaffected by CN variation. Duplications and deletions were primarily observed in the Threonine-rich region or Serine/Threonine-rich region located in the center or end of the *FLO5* gene, respectively. To better resolve intra-genic CN variation of *FLO11*, whose repeat unit is shorter than that of *FLO5*, we recalled CN variants with a smaller window size of 25 bp and re-evaluated CN variation (Figure S12). Using this window size, we found extensive duplications in 97 strains (73.5%) between gene coordinates 250-350 bp. Furthermore, duplications were observed in the hydrophobic Serine/Threonine-rich regions (Figure S12), which are associated with the flocculation phenotype (Fidalgo *et al*. 2006; Ramsook *et al*. 2010).

In contrast to *FLO5* and *FLO11*, other members of the *FLO* gene family did not exhibit intragenic CN variation. For example, CN variation in *FLO1* (*YAR050W*) and *FLO9* (*YAL063C*) typically spanned most or all of the sequence of each gene. Specifically, 125 strains (99.2%) had deletions spanning ≥80% of the gene in *FLO1* and only 2 strains (1.5%) had the entirety of the gene intact. *FLO9* had deletions in 99 strains (75%) that spanned ≥75% of the gene, 11 strains (8.3%) that had a partial deletion spanning <75% of the gene, whereas 1 strain (0.8%) had a CN of 2, and the remaining 21 strains (15.9%) had a CN of 1. In contrast, *FLO10* (*YKR102W*) showed limited CN variation. Specifically, 108 strains (81.8%) had no CN variation while 6 strains (4.5%) had deletions spanning the entirety of the gene. No duplications spanned the entirety of the gene but partial duplications were observed in 17 strains (12.9%) and were located in or just before the Serine/Threonine-rich region.

### Functional Implications CN Variable Genes

To determine the functional impact of deleted CN variable genes, we examined the relative growth of deleted CN variable genes (denoted with the Δ symbol) relative to the wild-type (WT) *S. cerevisiae* strain S288c across 418 conditions using the Hillenmeyer *et al*. 2008 data (Figure S13 and File S5). To determine the impact of duplicated genes, we examined growth fitness of the WT strain with low (~2-3 gene copies) or high plasmid CN (~8-24 gene copies), where each plasmid contained a single gene of interest from previously published data, relative to WT (Figure S14 and File S6; Payen *et al*. 2016).

Among deleted genes, 42 / 69 genes for which data exist showed negative and positive fitness effects in at least one tested condition in the S288c genetic background. Furthermore, we found that 12 / 42 genes that are commonly deleted among wine strains typically resulted in a fitness gain in conditions that resembled the fermentation environment. These conditions include growth at 23°C and at 25°C, temperatures within the 15-28°C range that wine is fermented in (Molina *et al*. 2007) and growth in minimal media, which is commonly used to understand fermentation-related processes (Seki *et al*. 1985; Govender *et al*. 2008; Vilela-Moura *et al*. 2008).

When examining fitness effects when grown at 23°C or at 25°C for 5 or 15 generations for the 12 commonly deleted genes, we observed at least one deletion that resulted in a fitness gain or loss for each condition. However, we observed extensive deletions in the *MAL1* locus (Figure 3) and therefore prioritized reporting the fitness impact of deletions in *MAL11*, *MAL12* and *MAL13*. *ΔMAL11* resulted in a fitness gain for growth at 23°C and 25°C for 5 (0.45X and 0.27X, respectively) and 15 generations (0.20X and 0.52X, respectively). *ΔMAL12* resulted in a gain of fitness at only 25°C after 15 generations (0.46X) and in a loss of fitness ranging from -0.36X to -1.29X in the other temperature conditions. Similarly, *ΔMAL13* resulted in fitness gains and losses dependent on the number of generations. For example, when grown for 15 generations at 25°C a fitness gain of 0.50X was observed while a fitness loss of -0.82X was observed at 23°C.

We next determined the fitness effect of deleted genes in minimal media after 0, 5, and 10 generations. Similar patterns of complex fitness gain and loss were observed as for the other conditions. For example, *ΔTHI12* resulted in a loss of fitness of -4.13X and -1.97X after 0 and 5 generations, but a fitness gain of 0.63X after 10 generations. In contrast, other genes resulted in positive fitness effects. For example, *ΔMAL12* resulted in a fitness gain of 7.25X and 10.41X for 0 generations and 10 generations.

Among duplicated genes, we focused on growth in glucose-and phosphate-limited conditions because glucose becomes scarce toward the end of fermentation prior to the diauxic shift and phosphate limitation is thought to contribute to stuck fermentations (Bisson 1999; Marsit and Dequin 2015). Among the 35 of the 69 genes where data were available, 14 genes had duplications among the 132 strains.

When examining fitness effects of duplicated genes in a glucose-limited environment in the S288c background, we found that fitness effects were small in magnitude and dependent on condition and plasmid CN (File S6). For example, *MAL32* low CN increased growth fitness by 0.02X but decreased fitness by -0.01X at a high CN (Figure S14). Interestingly, the most prevalent CN for *MAL32* across the 132 strains was 2 (96 strains, 72.7%), with only 3 strains showing a CN of 3 and none a higher CN. Another gene found at low CN in 37 strains (28%) was *HXT17*. Low plasmid CN in a glucose-limited conditions resulted in a fitness gain of 0.06X. In contrast, *MAL13* low or high plasmid CN resulted in a negative growth fitness of -0.02X and -0.01X, respectively. Interestingly, *MAL13* duplication is only observed in 4 strains (3%) and deletions are observed in 61 strains (46.2%).

Similar to the glucose-limited condition, we found fitness was dependent on high or low plasmid CN in the phosphate-limited condition. For example, *MAL31*, a gene present at low CN in 100 strains had a fitness gain of 0.04X at high plasmid CN but low plasmid CN resulted in a fitness loss of -0.02X. In contrast, *MAL32*, which was present at low CN in 99 strains, had a small fitness gain of 0.002X at low plasmid CN and a fitness loss at a high plasmid CN of -0.02X. A total of 6 genes resulted in a disadvantageous growth effect when present at low CN, such as *DDR48*, which resulted in a fitness loss of -0.04X. Altogether, our results suggest that the deleted and duplicated CN variable genes we observe (Figure 4) modulate cellular processes that result in advantageous fitness effects in conditions that resemble the fermentation environment.

**Figure 4.**
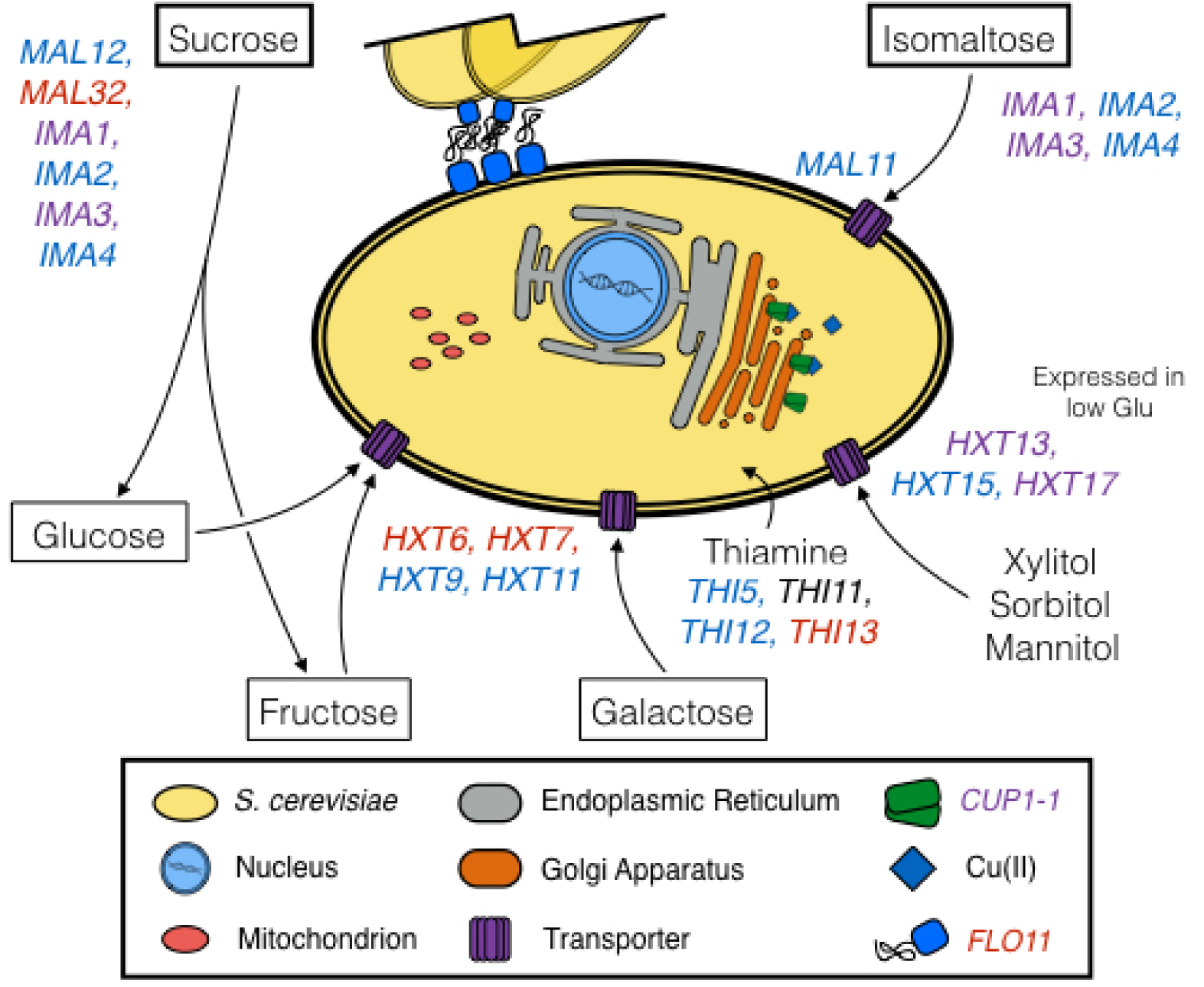
**Model summary of CN variable genes in wine yeast strains and their cellular functions.** Genes that are deleted among wine strains are blue whereas those that are duplicated are in red. Genes that were observed to be both duplicated and deleted (*IMA1, IMA3, HXT13, CUP-1*) are purple. Disaccharides are in thick-lined boxes, monosaccharides in thin-lined boxes, and alcohols are unboxed.

### Identifying loci absent from CN variation analysis

The present study was able to capture loci represented in the WT/S288c laboratory strain. To identify loci absent from the reference strain, we assembled unmapped reads for 20 strains with the lowest percentage of reads mapped and determined their identity (see methods; figure S4). Across the 20 strains, we identified 429 loci absent from S288c but present in other sequenced *S. cerevisiae* strains. These loci had an average length of 6.9 kb and an average coverage of 107.2X. The 20 loci with the highest bitscore alongside with the number of strains containing the locus are shown in Table S2. All but two of these loci were present only in one of the 20 strains we examined. The two exceptions were: the EC1118_1N26_0012p locus, which we found in 8 / 20 strains, which originates from horizontal gene transfer from *Zygosaccharomyces rouxii* to the commercial EC1118 wine strain of *S. cerevisiae* (Novo *et al*. 2009); and the EC1118_1O4_6656p locus, which we found in 7 / 20 strains. This locus was also originally found in the EC1118 strain (Novo *et al*. 2009) and contains a gene similar to a conserved hypothetical protein found in *S. cerevisiae* strain AWRI1631 (Borneman *et al*. 2008).

## Discussion

CN variant loci are known to contribute to the genomic and phenotypic diversity (Perry *et al*. 2007; Cutler and Kassner 2008; Orozco *et al*. 2009). However, the extent of CN variation in wine strains of *S. cerevisiae* and its impact on phenotypic variation remains less understood. Our examination of structural variation in 132 yeast strains representative of the ‘wine clade’ showed that CN variants are a significant contributor to the genomic diversity of wine strains of *S. cerevisiae*. Importantly, CN variant loci overlap with diverse genes and gene families functionally related to the fermentation environment such as *CUP, FLO, THI, MAL, IMA* and *HXT* (summarized in Figure 4).

The characteristics of CN variation in wine yeast (Figure 1A; Figure S6; Table S1) were found to be similar to those of the recently described beer yeast lineage (Gallone *et al*. 2016). For example, both lineages exhibited a similar size range of CNVRs (Figure 1A; Figure S6; Table S1) as well as a higher prevalence of CNVRs in the subtelomeric regions (Figure 1D). However, wine strains had a smaller fraction of their genome affected by CN variation (Figure S6) than beer strains (Gallone *et al*. 2016).

Wine yeast strains are thought to be partially domesticated due to the seasonal nature of wine-making, which allows for outcrossing with wild populations (Marsit and Dequin 2015; Gallone *et al*. 2016; Gonçalves *et al*. 2016). One human-driven signature of domestication is thought to be the duplication of the *CUP1* locus because multiple copies confer copper resistance and copper sulfates have been used to combat powdery mildews in vineyards since the early 1800s (Warringer *et al*. 2011; Marsit and Dequin 2015). Consistent with this ‘partial domestication’ view (Marsit and Dequin 2015; Gallone *et al*. 2016; Gonçalves *et al*. 2016), many wine strains were not CN variable for *CUP1-1* and *CUP1-2* or had one or both genes deleted (Figure 3).

An alternative, albeit not necessarily conflicting, hypothesis is that wine yeasts underwent domestication for specific but diverse wine flavor profiles (Hyma *et al*. 2011). Consistent with this view is the deletion (in >90% of the strains) of the *THI5* gene (Figure 3), whose activity is known to produce an undesirable rotten-egg sensory perception via higher SH_2_ production and is associated with sluggish fermentations (Bartra *et al*. 2010). In contrast to wine strains, duplications of *THI5* have been observed across the *Saccharomyces* genus, including in several strains of *S. cerevisiae* (CBS1171, 2 copies; S288c, 4 copies; EM93, 5 copies), *S. paradoxus* (5 copies), and the lager brewing yeast hybrid *Saccharomyces pastorianus* (syn. *S. carlsbergensis*; 2+ copies) (Wightman and Meacock 2003). In contrast*, THI13*, which is duplicated in 62.1% of strains, shows an increase in its expression 6-100-fold in *S. cerevisiae* when grown on medium containing low concentrations of thiamine allowing for the compensation of low thiamine levels (Li *et al*. 2010). Low levels of thiamine in wine fermentation have been associated with stuck or slow fermentations (Ough *et al*. 1989; Bataillon *et al*. 1996). Similar to *THI5* deletions, *THI13* duplications may have also been driven by human activity due to the advantageous effect of increased expression within the fermentation environment.

Two other gene families subject to CN variation were the *MAL* and *HXT* gene families. The S288c strain that we used as a reference contained two *MAL* loci (*MAL1* and *MAL3*), each containing three genes – a maltose permease (*MALx1*), a maltase (*MALx2*), and an MAL trans-activator (*MALx3*) – and located near the ends of different chromosomes (Michels *et al*. 1992). *MAL1* has been observed to be duplicated in beer strains of *S. cerevisiae* (Gallone *et al*. 2016; Gonçalves *et al*. 2016) while wine strains primarily lack this locus (Figure 3; Gonçalves *et al*. 2016). In contrast to the deletion of the *MAL1* locus, *MAL3* duplication in wine yeasts (Figure 3; Gonçalves *et al*. 2016) is surprising because maltose is absent from the grape must (Gallone *et al*. 2016). However, knockout studies have demonstrated *MAL32* is necessary for growth on turanose, maltotriose, and sucrose (Brown *et al*. 2010), which are present in small quantities in wines (Victoria and Carmen 2013). Due to the prominent duplication of *MAL3*, in particular the enzymatic genes *MAL31* and *MAL32*, we speculate that the *MAL3* locus may be utilized to obtain sugars less prevalent in the wine environment or serve other purposes.

The *HXT* gene family in the S288c strain that we used as a reference contains 16 *HXT paralogs, GAL2, SNF3* and *RGT2*. The expansion of the HXT gene family is positively correlated with aerobic fermentation in *S. paradoxus* and *S. cerevisiae* (Lin and Li 2011). *HXT6* and *HXT7* are high-affinity glucose transporters expressed at low glucose levels and repressed at high glucose levels (Reifenberger *et al*. 1995). In contrast to the recently described Asia (Sake), Britain (Beer) and Mosaic lineages (Gallone *et al*. 2016), we detected duplications in the *HXT6* and *HXT7* genes in wine yeasts (Figure 3). This may confer an advantage toward the end of fermentation and before the diauxic shift when glucose becomes a scarce resource. Evidence potentially supporting this hypothesis is that *HXT6* and *HXT7* are up-regulated by 9.8 and 5.6-fold, respectively, through wine fermentation in the *S. cerevisiae* strain Vin13 (Marks *et al*. 2008). Furthermore, *HXT6* or *HXT7* is found to be duplicated in experimentally evolved populations in glucose-limited environments (Dunham *et al*. 2002; Gresham *et al*. 2008; Dunn *et al*. 2012).

In summary, these results together with recent studies of CN variation in beer yeast strains (Gallone *et al*. 2016; Gonçalves *et al*. 2016), suggest that this type of variation significantly contributes to the genomic diversity of domesticated yeast strains. Furthermore, as most studies of CN variation, including ours, use reference strains, they are likely conservative in estimating the amount of CN variation present in populations. This caveat notwithstanding, examination of publically available data regarding the functional impact of duplicated or deleted genes (again in the context provided by the reference strain’s genetic background) suggests that CN variation in several, but not all, of the wine yeast genes confer fitness advantages in conditions that resemble the fermentation environment. Our results raise the questions of the extent to which CN variation contributes to fungal, and more generally microbial, domestication as well as whether the importance of CN variants in natural yeast populations, including those of other *Saccharomyces* yeasts, is on par to their importance in domestication environments.

## Acknowledgments

We thank members of the Rokas lab for helpful discussions and advice. This work was conducted in part using the resources of the Advanced Computing Center for Research and Education at Vanderbilt University. JS was supported by the Graduate Program in Biological Sciences at Vanderbilt University. This work was supported in part by the National Science Foundation (DEB-1442113 to A.R.).

